# Olfactory-Visual Integration Facilitates Scene Perception through Modulation of the Parahippocampal Place Area

**DOI:** 10.64898/2026.11.23.768575

**Authors:** Yanqing Wang, Kepu Chen, Yuli Wu, Wen Zhou

## Abstract

Visual scenes anchor our experience of the world, yet the places in which scenes unfold are defined not only by what we see but also by what we smell. Whether and how odors contribute to the neural representation of visual scenes remains poorly understood. Here we asked whether a semantically congruent odor facilitates scene perception and identified the neural mechanisms supporting this facilitation. In two behavioral experiments, participants judged whether briefly presented, masked images depicted intact or scrambled scenes while continuously smelling an odor. A congruent odor lowered the exposure duration needed for accurate scene detection, and a placebo experiment, in which participants merely believed they were smelling an odor, failed to reproduce this benefit, indicating that the facilitation depends on actual sensory input rather than semantic priming. In an fMRI experiment, congruent odors enhanced responses and sharpened category-level scene representations in the parahippocampal place area (PPA), with the congruency-driven enhancement localized to the same anterior subregion that encoded scene identity. A conjunction analysis revealed superadditive responses to congruent odor–scene pairs in the orbitofrontal cortex (OFC) and hippocampus, and psychophysiological interaction analyses showed that congruent stimulation strengthened functional coupling among the OFC, PPA, and piriform cortex. Notably, the piriform cortex decoded visual scene identity after congruent odor–scene pairing, suggesting that odor–scene binding extends scene representations into primary olfactory cortex. Together, these findings identify the PPA as a site where olfactory signals refine the neural code for visual scenes and reveal an OFC-coordinated piriform–OFC–PPA network through which smell shapes our sense of place.

## Introduction

Our perception of a scene extends beyond what is visible. A garden, a restaurant, or a street is encountered through a characteristic ensemble of sights, sounds, and smells, and it is this combination of inputs that makes a place feel distinct. Cognitive neuroscience has established a dedicated cortical network for scene processing that includes the parahippocampal place area (PPA), the occipital place area (OPA), and the retrosplenial cortex (RSC)(Malcolm et al., 2016). This network, however, has been characterized almost entirely on the basis of visual input. Although recent work has begun to ask how non-visual cues such as sounds contribute to scene representations(Häusler et al., 2022), whether olfactory cues are part of scene perception, and at what neural site such contributions might arise, remains unknown.

Odors hold a privileged relationship with places. Unlike most visual or auditory signals, an odor is rarely tied to a single object within a scene. It is diffuse and ambient, and it typically characterizes the environment as a whole. The aroma of bread does not point to a particular loaf but to the bakery itself. Humans appear to exploit this property in daily life. Olfaction guides navigation toward distant odor sources (Wu et al., 2020) and provides rich contextual information about the identity and quality of a place (Raithel & Gottfried, 2021). Odors are also potent triggers of vivid, emotionally toned autobiographical memories, often of the very settings in which they were first encountered(Arshamian et al., 2013). These observations suggest that smell may be an integral constituent of how scenes are perceived rather than an accessory signal.

Empirical work on olfactory-visual interactions has focused almost exclusively on discrete objects. Congruent odors facilitate the detection and identification of corresponding objects (Gottfried & Dolan, 2003), spontaneously draw visual attention toward congruent items (Chen et al., 2013; Fontana et al., 2025), and bias the perception of ambiguous visual stimuli in binocular rivalry (Zhou et al., 2010; Zhou et al., 2012). At the neural level, these interactions engage the orbitofrontal cortex (OFC) and hippocampus, which show enhanced responses to semantically congruent odor-picture pairs(Gottfried & Dolan, 2003). What this literature leaves open is whether odors also modulate the perception of complex visual scenes, which provide the coherent spatial context of daily experience, and if so, whether such modulation occurs within the cortical network dedicated to scene processing.

Among scene-selective regions, the PPA is the most likely site of such an interaction. It is critically involved in encoding and discriminating visual scenes(Baumann & Mattingley, 2016; Epstein et al., 1999; Mégevand et al., 2014; Walther et al., 2009; Walther et al., 2011). Its responses are not strictly visual, as it also processes spatial information conveyed through auditory scene descriptions (Häusler et al., 2022) and exhibits functional subdivisions tied to different aspects of scene content (Baldassano et al., 2016; Rolls et al., 2024). We therefore hypothesized that a semantically congruent odor would facilitate behavioral scene discrimination, refine scene representations within the PPA, and recruit a broader olfactory-visual network involving the piriform cortex, OFC, and hippocampus. To test these hypotheses, we conducted a series of experiments. Experiment 1 established the behavioral effect of olfactory-visual congruency on scene detection. Experiment 2 used a placebo manipulation to rule out a purely semantic priming account. Finally, fMRI combined with univariate, multivoxel pattern, and psychophysiological interaction analyses delineated the neural mechanisms underlying this crossmodal facilitation.

## Results

### Congruent olfactory-visual stimuli facilitate scene discrimination

In Experiment 1, we asked whether a semantically congruent odor facilitates visual scene discrimination. Participants were continuously exposed to an odor associated with either a rose garden or an auto shop while judging whether a rapidly presented image was an intact scene or a scrambled stimulus (Figure 1a). We analyzed scene detection accuracy using a 2 (olfactory-visual stimulus congruency, congruent vs. incongruent) × 7 (stimulus duration, 16, 25, 34, 43, 52, 61, and 70 ms) repeated-measures ANOVA. The analysis revealed a significant main effect of congruency (*F*_(1, 35)_ = 12.04, *p* = 0.001, *η*² = 0.256), which was driven by higher detection accuracy for congruent trials (mean ± SE: 0.81 ± 0.07) compared to incongruent trials (0.79 ± 0.07). As expected, the ANOVA also revealed a significant main effect of stimulus duration (*F*_(6, 210)_ = 315.08, *p* < 0.001, *η*² = 0.90) and a significant interaction between congruency and stimulus duration (*F*_(6, 210)_ = 2.48, *p* = 0.025, *η*² = 0.066). Simple effects analysis indicated that detection accuracy was significantly higher in the congruent condition than in the incongruent condition at the brief stimulus duration of 25 ms (*p* = 0.004). However, this difference was not observed at longer stimulus durations (*p*s > 0.05), likely due to performance approaching ceiling.

**Figure 1.**
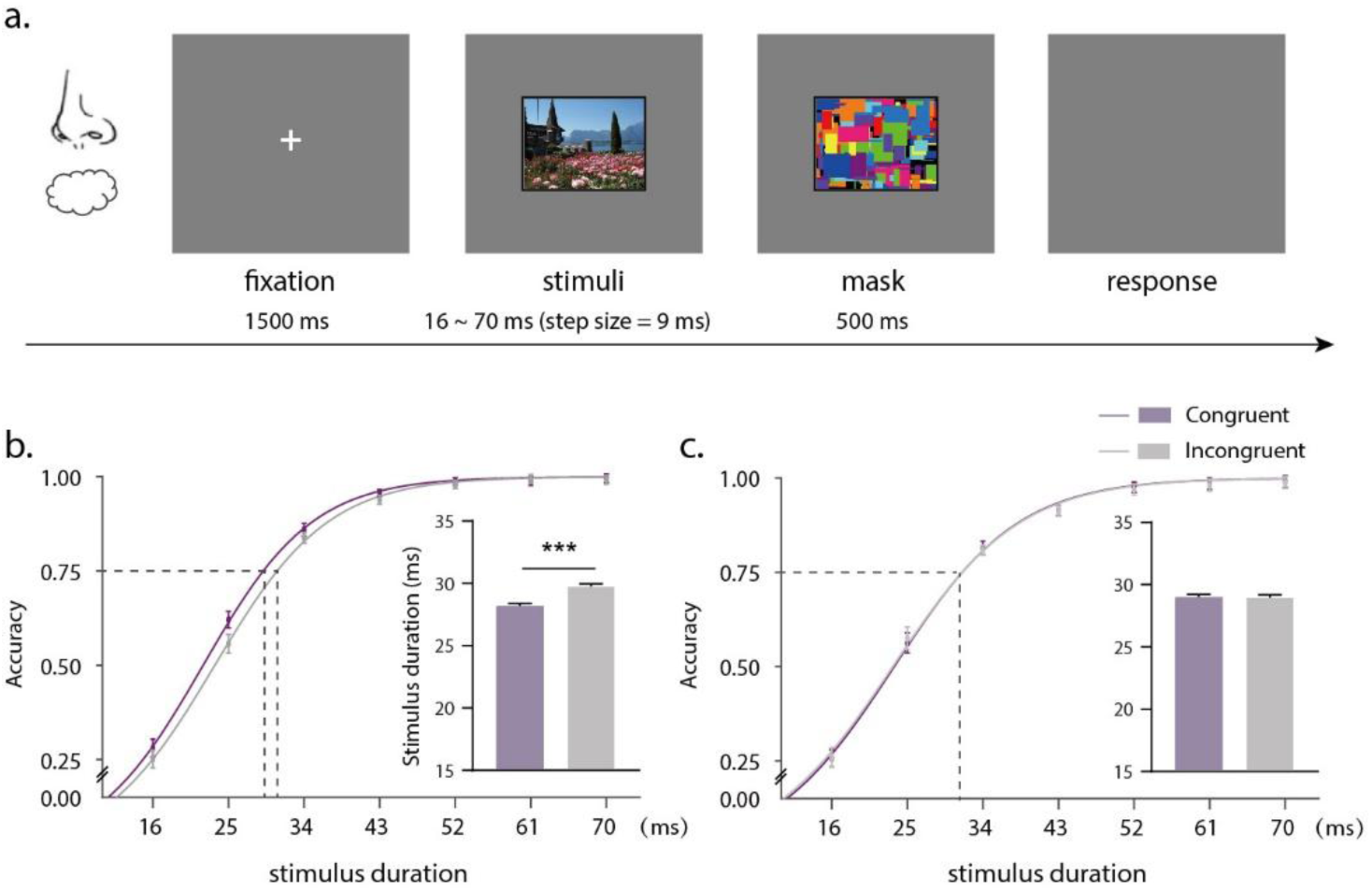
**a.** Experimental procedure and design of the task used in Experiments 1 and 2. **b.** Psychometric functions fitted to the accuracy data and the stimulus duration required to achieve 75% accuracy in Experiment 1. The presence of a congruent odor significantly reduced the threshold duration compared to the incongruent condition. **c.** Psychometric functions and the stimulus duration required to achieve 75% accuracy in Experiment 2. No significant difference was observed between conditions, ruling out a purely semantic priming account. Error bars represent standard errors of the mean adjusted for individual differences. *** : *p <* 0.001.

To complement the ANOVA, psychometric functions were calculated for each participant and condition by fitting a Boltzmann sigmoidal function to the scene detection accuracy data (Figure 1b). This analysis revealed that the stimulus duration required to achieve 75% accuracy was significantly reduced in the presence of a congruent odor compared to an incongruent odor (28.18 ms *vs.* 29.77 ms; *t*_(35)_ = 3.77, *p* = 0.001, Cohen’s *d* = 0.63). This finding indicates that participants exhibited enhanced scene discrimination sensitivity when smelling a congruent odor, even though the task did not require explicit scene discrimination.

While these results suggest a perceptual benefit of olfactory-visual congruency, it remains possible that the effect was driven by semantic priming, whereby the conceptual link between the odor (e.g., rose) and the scene (e.g., rose garden) facilitated visual recognition. In Experiment 2, we tested this possibility with a placebo manipulation. An independent sample of 36 participants performed the same scene detection task, but two bottles of purified water were used in place of the actual odors. Participants were deceived into believing that one bottle contained a low concentration of a rose-garden-related odor and the other contained a low concentration of an auto-shop-related odor, and the suggested odor label was displayed continuously at the start of each block. Post-experiment ratings confirmed that this manipulation successfully altered participants’ olfactory perceptions in the direction of the suggested odor content. When the water was suggested to contain a rose odor (compared to a gasoline odor), participants rated it as more rose-like (31.58 *vs.* 11.50; *t*_(35)_ = 7.03, *p* < 0.001), less gasoline-like (13.5 *vs*. 30.03; *t*_(35)_ = 5.02, *p* < 0.001), more pleasant (41.19 *vs.* 31.33; *t*_(35)_ = 3.30, *p* = 0.002), and weaker in intensity (23.03 *vs*. 28.25; *t*_(35)_ = 2.12, *p* = 0.041). Crucially, despite this successful psychological manipulation, scene detection accuracy remained unaffected. A repeated-measures ANOVA revealed a significant main effect of stimulus duration (*F*_(6, 210)_ = 271.80, *p* < 0.001, *η*² = 0.87), but neither the main effect of congruency (*F*_(1, 35)_ = 1.52, *p* = 0.23, *η*² = 0.042) nor the interaction between congruency and stimulus duration (*F*_(6, 210)_ = 0.69, *p* = 0.655, *η*² = 0.019) reached significance. Furthermore, the psychometric curves for the congruent and incongruent conditions overlapped entirely, and the stimulus duration yielding 75% accuracy did not differ significantly between conditions (*t*_(35)_ = 0.068, *p* = 0.946; Figure 1c). These findings confirm that the facilitation depends on sensory congruency between actual olfactory and visual inputs rather than on semantic priming.

### Congruent odors enhance scene-evoked activity in the PPA

In the fMRI experiment, we asked where congruent odors modulate the neural processing of visual scenes. Participants performed an odor detection task while scenes and odors were presented. On each trial they indicated with a button press whether an odor was present, and they performed this task with high accuracy (0.90 ± 0.01), confirming reliable task engagement. Because the task required no judgment about the scenes, any modulation of scene-related activity by odor congruency would reflect automatic crossmodal influence rather than task-dependent processing. To localize the sites of such modulation, we compared activity evoked by the bimodal congruent (OVc) and incongruent (OVi) conditions. The congruent condition evoked significantly greater activation in the bilateral parahippocampal place area (left PPA, x = −32, y = −36, z = −18; right PPA, x = 38, y = −40, z = −14; Figure 2b). A region-of-interest (ROI) analysis extracting the mean parameter estimates from the bilateral PPA confirmed a significantly greater response to the OVc condition compared to the OVi condition in both the left PPA (*t*_(30)_ = 10.00, *p* < 0.001) and the right PPA (*t*_(30)_ = 7.89, *p* < 0.001).

**Figure 2.**
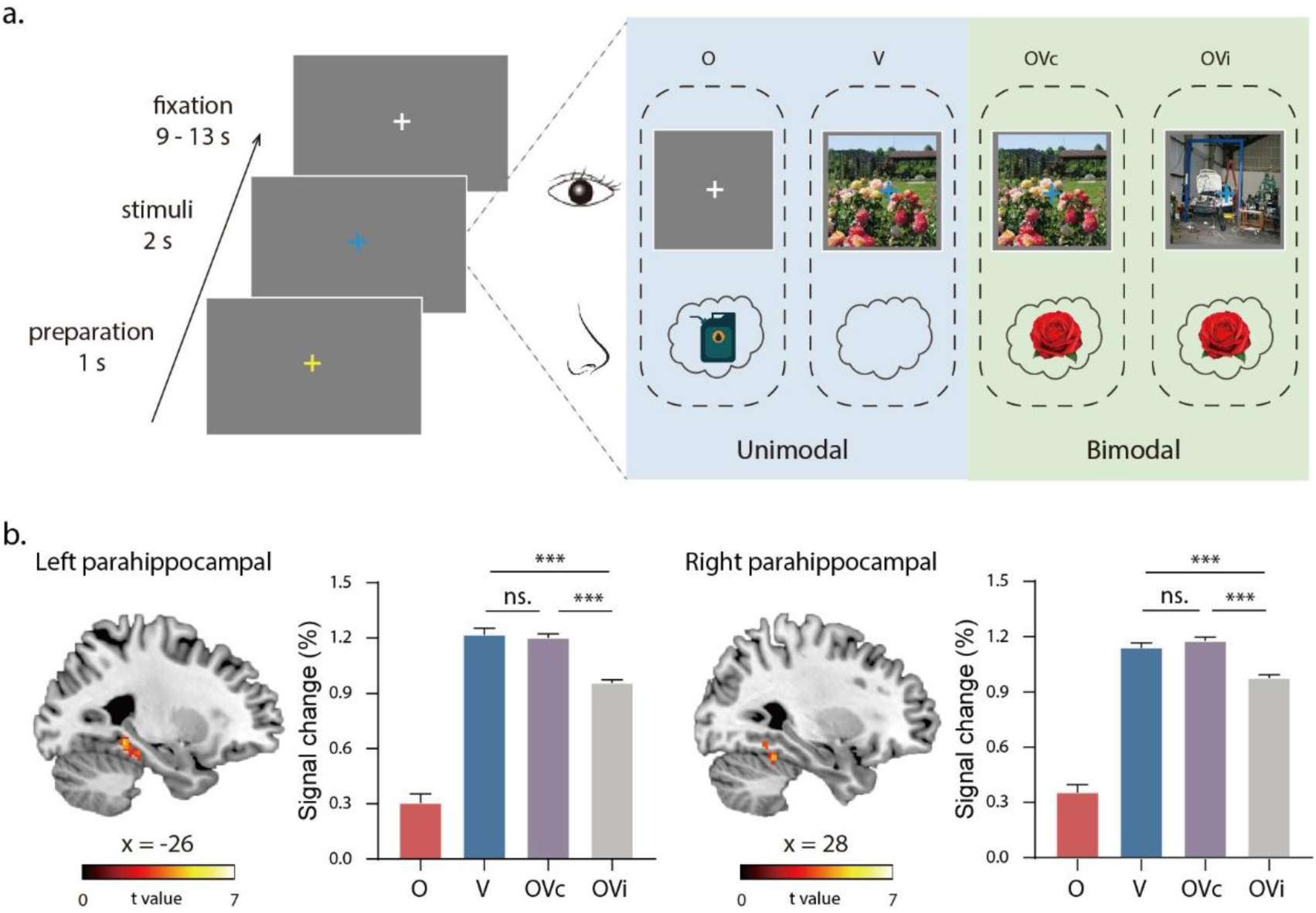
**a.** Procedure of the odor detection task used in the fMRI experiment. Each trial began with a 1-s yellow fixation cross, indicating imminent stimulus onset and signaling participants to pause breathing and prepare to sniff. The fixation cross then turned blue, alerting participants to the start of a new breathing cycle. As participants inhaled, stimuli were presented for 2000 ms. Four types of trials were included: (1) unimodal olfactory (O) trials, in which only an odor was presented; (2) unimodal visual (V) trials, in which only a visual image was presented; (3) bimodal congruent (OVc) trials, in which olfactory and visual stimuli were presented simultaneously and carried matching semantic information (e.g., rose odor paired with a rose garden image); and (4) bimodal incongruent (OVi) trials, in which olfactory and visual stimuli were presented simultaneously but carried mismatched information (e.g., rose odor paired with an auto shop image). **b.** Regions showing enhanced activation during the OVc > OVi contrast, including the bilateral parahippocampal place area (PPA), and the estimated percentage signal change within the PPA.

### Congruent odors sharpen scene representations in the PPA

Enhanced activity does not by itself indicate that the neural code for scenes has changed. We therefore next asked whether congruent odors refine the representational content of scene-selective cortex, using a two-stage multivoxel pattern analysis (MVPA) approach in which voxel clusters identified by a whole-brain searchlight were subsequently analyzed as independent ROIs. The searchlight analysis first identified regions encoding scene category. As illustrated in Figure 3a, decoding accuracy was significantly above chance in large clusters within the bilateral PPA (x = −14, y = −56, z = −8; x = 30, y = −46, z = −10), the RSC (x = −8, y = −62, z = 12; x = 16, y = −60, z = 0), and the left OPA (x = −34, y = −80, z = 26), consistent with previous work on scene encoding and discrimination. ROI-based classification analyses then compared decoding accuracy across the unimodal visual (V), OVc, and OVi conditions. These analyses revealed a significant main effect of condition in the PPA (*F*_(2, 60)_ = 3.63, *p* = 0.03, *η*² = 0.108; Figure 3c). Post hoc tests showed significantly higher classification accuracy for the OVc condition compared to both the V condition (*t*_(30)_ = 2.30, *p* = 0.03) and the OVi condition ((*t*_(30)_ = 2.37, *p* = 0.02), with no difference between the OVi and V conditions (*t*_(30)_ = 0.29, *p* = 0.78). In contrast, no main effect of condition was observed in the OPA (*F*_(2, 36)_ = 0.57, *p* = 0.57, *η*² = 0.03) or the RSC (*F*_(2, 60)_ = 0.01, *p* = 0.99, *η*² < 0.001). These findings indicate that congruent odors sharpen scene representations selectively within the PPA.

**Figure 3.**
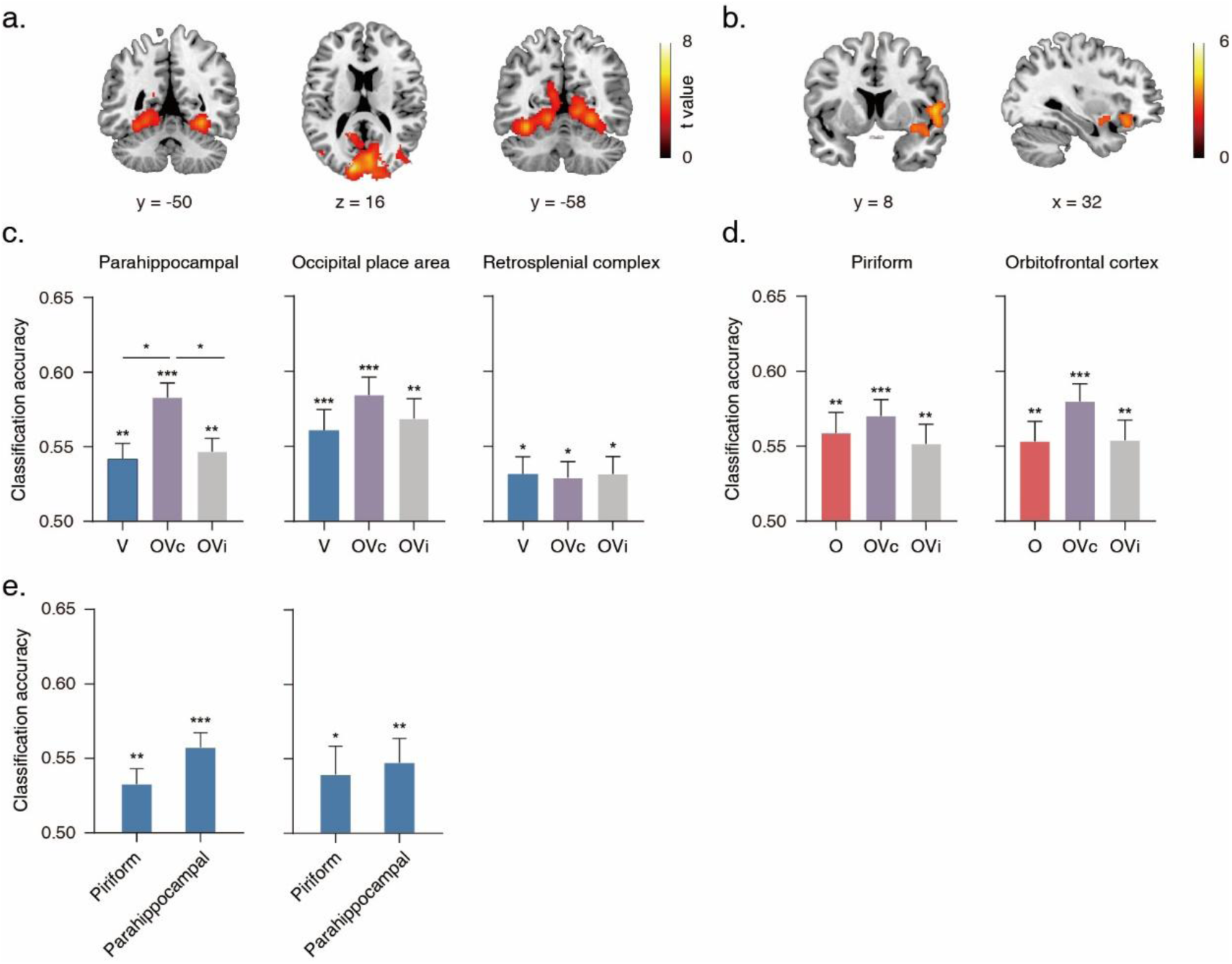
**a.** Regions from which whole-brain searchlight MVPA could decode whether the visual stimulus was an auto shop or a rose garden at above-chance accuracy. **b.** Regions from which whole-brain searchlight MVPA could decode whether the olfactory stimulus was rose or gasoline at above-chance accuracy. **c.** Classification accuracy in the PPA, OPA, and RSC for decoding the visual stimulus in the V, OVc, and OVi conditions. **d.** Classification accuracy in the piriform cortex and OFC for decoding the olfactory stimulus in the O, OVc, and OVi conditions. **e.** ROI-based MVPA results for visual scene decoding in the piriform cortex and parahippocampal place area (PPA). The left panel shows that both regions could significantly discriminate between rose garden and auto shop scenes when using all visual-containing trials (V, OVc, and OVi conditions). The right panel shows that classification accuracy remained significantly above chance even when training and testing the classifier exclusively on unimodal visual (V) trials. Error bars represent standard errors of the mean adjusted for individual differences. * : *p* < 0.05; ** : *p* < 0.01; *** : *p* < 0.001.

Several recent studies have proposed functional subdivisions within the PPA (Baldassano et al., 2016; Rolls et al., 2024). To test whether the voxels encoding scene category overlap with those enhanced by olfactory-visual congruency, we generated two spatial probability maps, one quantifying, for each PPA voxel, the proportion of participants with above-chance scene classification, and the other quantifying the proportion of participants showing a greater response to OVc than to OVi. As illustrated in Figure 4, the peak voxels of both maps were located in the anterior PPA, demonstrating strong spatial consistency. This finding suggests that the specific subset of PPA voxels that represents scene information is selectively enhanced by congruent olfactory cues.

**Figure 4.**
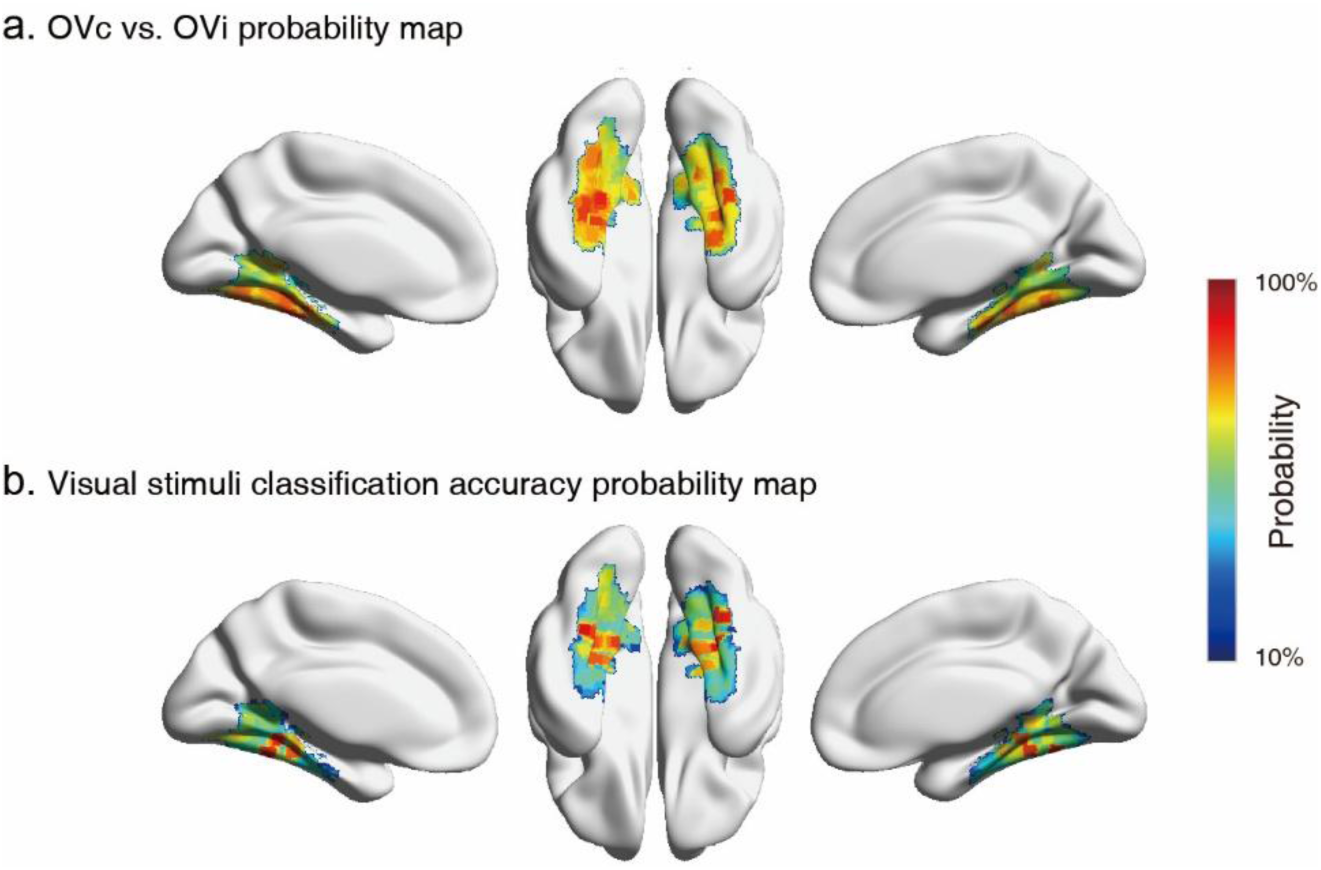
**a.** Group-level probability maps illustrating the voxel-wise proportion of participants who showed statistically enhanced activation for the contrast OVc > OVi. **b.** Group-level probability maps illustrating the voxel-wise proportion of participants exhibiting above-chance decoding accuracy when classifying the visual stimulus category (auto shop vs. rose garden).

### Odor category coding remains stable while scene information extends into the piriform cortex

A separate question concerns the other side of the pairing, namely how odor representations are affected by concurrent scene processing. To examine this, we compared classification accuracies for the unimodal olfactory (O), OVc, and OVi conditions in olfactory regions, mirroring the approach used for the visual scene decoder. Searchlight results revealed robust odor category coding in the right piriform cortex (x = 32, y = 2, z = −16) and the OFC (x = 30, y = 28, z = −14; Figure 3b). ROI-based classification analyses, however, did not reveal a main effect of condition in either the piriform cortex (*F*_(2, 60)_ = 0.57, *p* = 0.69, *η*² = 0.01) or the OFC (*F*_(2, 60)_ = 0.93, *p* = 0.40, *η*² = 0.03, Figure 3d), indicating that odor category coding in these regions remained stable across conditions.

Previous findings indicate that piriform cortex neurons carry a robust spatial representation of the environment (Poo et al., 2022). We therefore tested whether the piriform cortex encodes visual scenes following scene-odor binding. ROI-based classification analyses discriminating piriform activity patterns evoked by rose garden versus auto shop scenes revealed classification accuracy significantly above chance (*t*_(30)_ = 3.19, *p* = 0.003, Figure 3e left). Notably, this accuracy remained above chance even when the classifier was trained and tested exclusively on the unimodal visual condition (*t*_(30)_ = 2.08, *p* = 0.046, Figure 3e right). This crossmodal decoding suggests that scene-odor binding enables the piriform cortex to represent visual scene information.

### OFC and hippocampus integrate congruent odor-scene information

The results so far show that congruent odors modulate both the activity and the representational content of the PPA. We next asked which regions bind olfactory and visual scene inputs into an integrated representation. To this end, we conducted a conjunction analysis identifying regions that responded to both unimodal inputs and exhibited enhanced responses to bimodal presentations. This superadditive effect is a hallmark of multisensory integration. Significant clusters were found in the bilateral OFC and the hippocampus (Figure 5a). One-sample t tests confirmed that all three regions responded significantly to the O, V, OVc, and OVi conditions (all *t*(30) > 2.57, *p* < 0.015). A one-way repeated-measures ANOVA revealed a significant main effect of condition in the left OFC (*F*_(3, 90)_ = 35.40, *p* < 0.001), the right OFC (*F*_(3, 90)_ = 48.48, *p* < 0.001), and the hippocampus (*F*_(3, 90)_ = 47.43, *p* < 0.001). Post hoc tests revealed three key patterns. First, all three regions exhibited significantly greater activation for the OVc condition compared to either the O or V conditions (all *p* < 0.001). Second, activation did not differ between the O and V conditions (all *p* > 0.13). Third, the OVi condition also elicited stronger activation than either unimodal condition, showing greater activation than the O condition (all *p* < 0.005) and the V condition (all *p* < 0.004). Critically, all three regions showed significantly greater responses to the OVc condition than to the OVi condition (all *p* < 0.001). This specific enhancement for congruent olfactory-visual stimulation underscores the role of the OFC and hippocampus in binding crossmodal scene features into a coherent percept.

**Figure 5.**
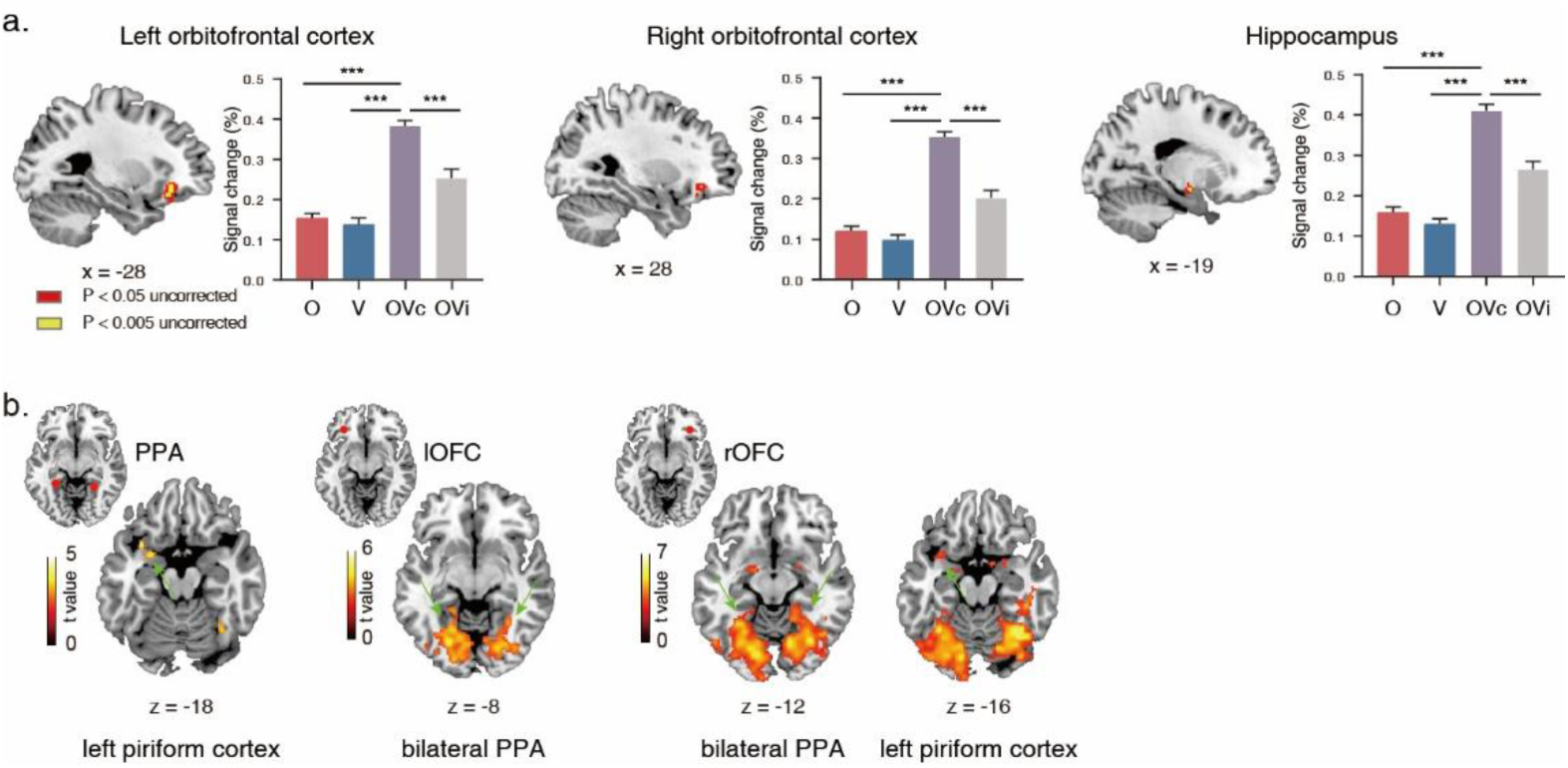
**a.** Regions identified by the conjunction analysis revealing olfactory-visual scene integration in the bilateral orbitofrontal cortex (OFC) and hippocampus, along with the estimated percentage signal change for each condition in these regions. Voxels in red are significant at *p* < 0.05 uncorrected, whereas voxels in yellow remain significant at *p* < 0.005 uncorrected. **b.** Regions showing increased functional connectivity with the PPA, left OFC (lOFC), and right OFC (rOFC) for the contrast OVc > OVi. The results are displayed at an uncorrected cluster-level threshold of *p* < 0.001.

### Congruent stimulation strengthens functional coupling within a piriform-OFC-PPA network

The final question is how the congruency effects in the PPA and the integrative responses in the OFC and hippocampus are coordinated. We reasoned that olfactory modulation of the PPA might be mediated by altered functional connectivity between the PPA and olfactory regions, and we tested this hypothesis with psychophysiological interaction (PPI) analyses using the PPA, left OFC, right OFC, and hippocampus as seed regions, each defined individually for every participant. The PPI analysis with the PPA seed revealed increased functional coupling with the left piriform cortex (x = −26, y = 2, z = −16; *p*_FWE_ < 0.05, small-volume corrected [SVC], Figure 5b left). The right OFC seed showed enhanced connectivity with the left piriform cortex (x = −32, y = 10, z = −16; *p*_FWE_ < 0.05, SVC) and the bilateral PPA (x = −28, y = −52, z = −10; x = 16, y = −40, z = −6, Figure 5b right). Similarly, the left OFC seed showed enhanced connectivity with the bilateral PPA (x = −20, y = −58, z = −8; x = 30, y = −42, z = −16, Figure 5b middle). In contrast, the PPI analysis with the hippocampal seed revealed no significant connectivity changes for the congruent relative to the incongruent condition at the corrected threshold. This pattern of results suggests a functional division within the integration network. The OFC, but not the hippocampus, showed congruency-dependent changes in coupling with both unimodal regions, consistent with its proposed role as the central coordinator of olfactory-visual integration, whereas the hippocampus appears to contribute integrative responses without exhibiting detectable modulation of its connectivity. These findings delineate a distributed network in which the OFC coordinates the exchange of olfactory and visual inputs, thereby supporting the integration of sensory cues into a coherent scene percept.

## Discussion

The present study asked whether odors are part of scene perception and identified the neural mechanisms through which they contribute to it. We found that a semantically congruent odor improved the accuracy and perceptual sensitivity of scene discrimination, and a placebo control confirmed that this effect depends on actual sensory congruency rather than on abstract semantic priming. At the neural level, the parahippocampal place area (PPA), a key region of the visual scene-processing network, emerged as a primary locus of this crossmodal interaction, showing both enhanced activation and sharpened scene representations in the presence of a congruent odor. The orbitofrontal cortex (OFC) and hippocampus exhibited superadditive responses to congruent odor-scene pairs, and congruent stimulation strengthened functional coupling within a piriform-OFC-PPA network. Together, these findings identify a specific mechanism whereby olfactory signals refine the neural code for visual scenes and delineate the network that supports this integration.

### Functional specificity of the PPA in olfactory-visual integration

A central finding of our study is the robust modulation of the PPA by congruent olfactory information. The PPA is a cornerstone of the cortical network for scene perception, playing a critical role in recognizing individual scenes and ascertaining their spatial structure(Epstein & Baker, 2019). Direct electrical stimulation of this region evokes hallucinations of visually experienced places(Mégevand et al., 2014), and lesions involving the PPA impair scene recognition, establishing its causal contribution to scene perception. In all of this work, however, the manipulations were exclusively visual, so whether the inputs driving the PPA are confined to vision has never been directly tested. Our results speak to this untested assumption. PPA activity was enhanced for congruent olfactory-visual pairs, and the multivoxel patterns representing scene categories were significantly sharpened by a congruent odor. This representational enhancement indicates that olfaction does not merely provide a generic contextual boost but actively refines the neural code for visual scenes within the PPA.

This finding extends a growing body of evidence suggesting that the PPA integrates information beyond the visual modality. The PPA has been shown to respond to spatial information conveyed through auditory descriptions of scenes, indicating a capacity for amodal spatial processing(Häusler et al., 2022). Our results provide the first direct evidence for olfactory modulation of scene processing in the PPA. The spatial overlap we observed within the anterior PPA between voxels that discriminate scene category and those that show congruency-based enhancement further supports the idea that a specific sub-region of the PPA serves as a direct site of this crossmodal interaction. This aligns with the view that the PPA is not a monolithic entity but possesses functional subdivisions (Baldassano et al., 2016; Rolls et al., 2024).

Notably, we did not observe similar olfactory modulation in the other two canonical scene-selective regions, the retrosplenial cortex (RSC) and the occipital place area (OPA). While all three regions were involved in encoding visual scene identity, the specificity of the olfactory effect to the PPA suggests a functional dissociation within the scene-processing network. The PPA is thought to be more concerned with the representation of the local visual scene and its contents (Epstein, 2008; Park et al., 2011), whereas the RSC is more involved in situating the scene within a broader spatial environment for navigation (Park & Chun, 2009). The OPA, which is causally involved in scene perception (Dilks et al., 2013), has been specifically implicated in representing navigational affordances and ego-motion (Kamps et al., 2025). It is plausible that ambient olfactory cues, which provide rich contextual information about the identity and quality of a place, are more relevant for modulating the representation of the local scene’s identity in the PPA than for the navigational or broader spatial mapping functions of the OPA and RSC.

### A dynamic network orchestrating olfactory-visual integration

Our study moves beyond the identification of a single locus of integration by delineating a functional network that orchestrates this process. The superadditive responses observed in the OFC and hippocampus represent a hallmark of multisensory integration, confirming their roles as higher-order convergence zones. The OFC is a well-established hub for olfactory-visual integration, particularly for processing the affective and identity dimensions of congruent stimuli (Gottfried & Dolan, 2003). The hippocampus, crucial for associative memory, likely mediates the binding of olfactory and visual features into a coherent, contextually rich memory trace (Gottfried & Dolan, 2003). Our results extend the function of this network into the domain of complex scene perception.

Crucially, our psychophysiological interaction analyses illuminate the dynamic interplay between these regions. During congruent stimulation, functional connectivity increased between the OFC and both the PPA and the piriform cortex. The OFC thus appears to act as a central coordinator that receives convergent sensory inputs, evaluates their semantic congruency, and then, via top-down modulation, enhances processing in lower-level unimodal areas. The strengthened coupling with the PPA likely facilitates the refinement of the visual scene representation, while enhanced coupling with the piriform cortex may reflect a feedback mechanism that sharpens the ongoing olfactory representation. Consistent with this coordinator account, the hippocampus, despite its superadditive responses, did not show congruency-dependent changes in coupling with unimodal regions, suggesting that its contribution reflects the integration of already-converged signals rather than the coordination of processing across modalities. This model proposes a dynamic, tripartite network comprising the piriform cortex, OFC, and PPA that underlies the behavioral facilitation we observed.

An intriguing and unexpected finding was that the piriform cortex, the primary cortical recipient of olfactory information, could be used to successfully decode visual scene identity. This crossmodal decoding was successful even when the classifier was trained and tested exclusively on unimodal visual trials, indicating that after associative pairing, the piriform cortex can represent visual information. This suggests a more profound level of sensory integration than previously thought, whereby primary sensory areas may acquire representations of associated stimuli from other modalities, blurring the boundary between unimodal and multimodal cortex.

### Behavioral facilitation and the correspondence between the behavioral and fMRI paradigms

The behavioral facilitation observed in our study is consistent with the broader principle that congruent multisensory information enhances perceptual processing (Stein & Stanford, 2008). The effect was most pronounced at very brief stimulus durations, suggesting a rapid, automatic integration process operating at early stages of perception rather than a slower, deliberate cognitive strategy. A key methodological strength of our study is the placebo control experiment, which rules out an account based purely on abstract semantic priming. By demonstrating that the mere belief of smelling a congruent odor was insufficient to produce the behavioral effect, we confirmed that the facilitation arises from the brain’s integration of actual, concurrent sensory inputs.

The behavioral experiments used brief masked presentation, whereas the fMRI experiment presented scenes for 2000 ms, and this difference merits explanation. The two paradigms were designed to answer different questions. In the behavioral experiments, detection accuracy approached ceiling at longer durations, which left little room for facilitation to be expressed, and stimulus duration was therefore varied to place performance within a sensitive range. The short presentation times were the condition under which the behavioral effect is observable, not the condition under which it exists. The fMRI experiment addressed a complementary question, namely whether congruent odors modulate scene processing in the brain when scene discrimination is not required and performance cannot index the effect. Because the congruent and incongruent conditions in the scanner were matched in presentation duration, the enhanced activity and sharpened representations we observed in the PPA are attributable to stimulus congruency rather than to exposure duration. Furthermore, stimuli presented for only tens of milliseconds evoke hemodynamic responses that are difficult to detect reliably with BOLD fMRI, whereas the 2000 ms presentation ensured robust signal-to-noise for univariate and multivoxel analyses. The congruency effects observed under prolonged viewing therefore indicate that olfactory modulation of scene processing does not depend on brief exposure and persists when scenes are processed incidentally, a property consistent with the automatic, attention-independent character suggested by the behavioral data. In this respect, the two paradigms are complementary, with the behavioral experiments establishing that the effect is perceptually consequential under near-threshold conditions, and the fMRI experiment revealing that its neural signature is present and robust even when the perceptual benefit itself is not measurable.

The present findings may also speak to the well-documented power of odors to evoke vivid and emotionally rich memories of past places(Arshamian et al., 2013). Our results suggest one possible mechanism for this phenomenon. If a congruent odor sharpens the neural representation of a scene at the moment of encoding, as observed in the PPA, the resulting memory trace may itself be richer and more distinctive. Enhanced perception at encoding could thus constitute an early step toward the especially vivid recollections that odor-cued memories are known to produce. This account remains speculative, as we did not assess memory directly, but it generates a testable prediction that odor-cued enhancement of scene processing should strengthen subsequent scene memory.

### Limitations and future directions

While this study provides novel insights, several limitations highlight avenues for future research. First, our use of static scenes and continuous odor delivery limits ecological validity. Future studies employing virtual reality could investigate olfactory-visual integration during active navigation, capturing the dynamic sensory changes characteristic of real-world experience. Second, our fMRI task was designed to probe automatic integration by directing attention away from the scenes. It remains an open question how top-down attention would modulate the network we identified. An explicit task requiring participants to judge the congruency of odor-scene pairs might reveal stronger or more widespread modulation, potentially involving prefrontal control networks. Third, the limited temporal resolution of fMRI obscures the precise dynamics within the piriform-OFC-PPA network. Future research using MEG or EEG could map the temporal flow of information, determining whether olfactory input modulates visual processing via early feedforward sweeps or later feedback signals. Finally, the link between congruency-driven enhancement at encoding and subsequent memory remains to be tested empirically in future studies.

## Conclusion

In summary, this study reveals a specific neural mechanism whereby olfaction enhances visual scene perception. We demonstrate that congruent odors facilitate scene discrimination by refining neural representations within the PPA. This modulation is orchestrated by a broader network anchored by the OFC and hippocampus, which integrates multisensory inputs and coordinates processing across unimodal cortices via dynamic changes in functional connectivity. These findings significantly advance our understanding of both scene perception and multisensory integration, challenging the traditional view of the scene-selective cortex as a purely visual processor. By showing how the brain seamlessly synthesizes information across multiple senses, this work underscores the fundamentally multisensory nature of our perceived reality and the profound sense of place.

## Materials and Methods

### Participants

A total of 110 healthy non-smokers participated in the study. The sample included 36 individuals in Experiment 1 (15 males, mean age ± SD = 20.94 ± 2.79 years), 36 individuals in Experiment 2 (17 males, 22.58 ± 1.18 years), and 38 individuals in the fMRI experiment (13 males, 24.76 ± 2.35 years). All participants had normal olfactory function and normal or corrected to normal vision. Furthermore, they reported no current nasal obstruction, respiratory allergies, or upper respiratory infections at the time of testing. The study protocol was approved by the Institutional Review Board of the Institute of Psychology at the Chinese Academy of Sciences. Written informed consent was obtained from all participants prior to their involvement in the experiments.

For the fMRI experiment, data from seven participants were excluded from further analysis due to excessive head motion during the scan (i.e., > 3 mm) or inadequate task performance (i.e., < 75% accuracy). Consequently, data from the remaining 31 participants were included in the final fMRI analyses, ensuring the reliability and quality of the neuroimaging results.

### Stimuli

#### Visual stimuli

The visual stimuli consisted of color images downloaded from the internet, featuring two distinct categories of rose gardens and auto shops with 24 images per category. For the fMRI experiment, half of the images from each category were used for the experimental runs, while the remaining half served as localizer runs. For Experiments 1 and 2, the visual stimuli included the 24 intact images from each category, as well as 48 scrambled versions of the rose garden and auto shop images. All pictures were presented on a computer screen and matched in size.

#### Olfactory stimuli

The olfactory stimuli differed across experiments. In Experiment 1, the stimuli consisted of a rose odor created with rose oil at 2% v/v in propylene glycol and a gasoline odor created with gasoline at 0.1% v/v in propylene glycol. In the fMRI experiment, higher concentrations were used to ensure reliable detection inside the scanner, specifically rose oil at 80% v/v and gasoline at 5% v/v in propylene glycol. In Experiment 2, two bottles of purified water served as placebo controls for Experiment 1. Except for the water, all odors were suprathreshold. The odors were selected such that each was semantically congruent with one visual category, pairing rose with rose garden and gasoline with auto shop, and incongruent with the other. In Experiments 1 and 2, odors were presented using identical 40 ml polypropylene jars. Each jar contained 10 ml of clear liquid and was connected to two Teflon nosepieces via a Y shaped structure. Participants were instructed to insert the nosepieces into their nostrils and to continuously inhale through the nose and exhale through the mouth. In the fMRI experiment, odors were delivered via an olfactometer controlled by the stimulus computer.

Prior to Experiment 1, an independent sample of 26 participants (13 males; mean age ± SD = 22.65 ± 1.16 years) performed an odor rating task. They continuously inhaled either the rose or gasoline odor for 1 minute, repeated four times. After the first inhalation, they rated the odor intensity, pleasantness, and similarity to its corresponding visual stimulus on a 100 unit visual analog scale (VAS). After the second, third, and fourth inhalations, they rated only the intensity. A 1 minute break was provided between the two odors. After the first inhalation, gasoline was rated as more unpleasant compared to rose odor, with mean ratings of 25.27 versus 73.84 (*t*_(25)_ = 10.24, *p* < 0.001), and more intense, with mean ratings of 83.42 versus 77.62 (*t*_(25)_ = 2.28, *p* = 0.031). However, due to rapid adaptation to the gasoline odor, perceived intensities became comparable between the two odors during the second through fourth inhalation (77.15 vs. 74.08, 72.15 vs. 70.08, 65.50 vs. 65.23; all *p* > 0.05). Similarly, 22 participants from the fMRI experiment sampled each odor with a 1 minute break in between and rated its intensity, pleasantness, and similarity to the corresponding visual stimulus on a 7 point Likert scale. The results confirmed that the gasoline odor was perceived as more unpleasant than the rose odor, with mean ratings of 2.59 versus 4.68 (*t*_(21)_ = 6.50, *p* < 0.001), and more intense, with mean ratings of 5.45 versus 4.68 (*t*_(21)_ = 3.93, *p* = 0.001). Overall, the identification and rating tasks confirmed that the odors used in the present study were easily identifiable.

### Respiratory monitoring

Respiration was monitored continuously during fMRI scanning using a BioPac MP150 system and accompanying AcqKnowledge software from BioPac Systems. Breathing belts were placed around the chest and abdomen of each participant to accurately record respiratory patterns. For each experimental condition, respiratory features including inhalation duration and volume were calculated using the BreathMetrics toolbox. A repeated measures ANOVA with four conditions (O, V, OVc, and OVi) was conducted to compare these respiratory parameters. The analysis found no significant differences in inhalation duration (*F*_(3, 90)_ = 1.73, *p* = 0.17) or inhalation volume (*F*_(3, 90)_ = 0.75, *p* = 0.52) across conditions. These results confirm that the observed neural and behavioral effects cannot be attributed to differences in breathing patterns across the experimental conditions.

### Experimental design

#### Behavioral experiment

Visual stimuli were presented on a 24.5 inch LCD monitor with a 240 Hz refresh rate. Each trial began with a central fixation cross for 1500 ms followed by a visual stimulus presented for one of seven durations (16, 25, 34, 43, 52, 61, or 70 ms). Immediately at the offset of the visual stimulus, dynamic noise patches were presented for 500 ms to create a backward masking effect and precisely control the effective presentation duration of the visual stimuli (Figure 1a). Participants were then instructed to press the left or right arrow key as accurately as possible to indicate whether the presented image was an intact scene or a scrambled stimulus. Participants were continuously exposed to one olfactory stimulus per block and were given a break of at least 1 minute between blocks to prevent olfactory habituation. Each participant completed eight blocks of 70 trials each, with four blocks per olfactory condition (rose odor vs. gasoline odor). For each olfactory condition, a total of 280 trials (10 trials × 7 stimuli durations × 4 visual stimuli types) were randomly presented across the four blocks. Consequently, for each stimulus duration, there were 20 congruent olfactory visual events and 20 incongruent olfactory visual events presented throughout the experiment. Before the experimental blocks, each participant completed a practice block of 70 trials to become familiar with the task rules. In Experiment 1, actual rose or gasoline odors were presented to participants. In Experiment 2, participants were presented with two bottles of purified water but were deceived into believing they contained low concentrations of rose and gasoline odors, respectively. Before each block, participants were informed which odor they would receive and subsequently rated the samples using a 100-unit VAS for similarity to rose/gasoline, intensity, and pleasantness. Furthermore, the suggested odor label was displayed continuously on the central screen at the start of each block, acting as a salient semantic cue.

Detection accuracies were submitted to a two way repeated measures ANOVA with factors of congruency (congruent versus incongruent) and stimulus duration (16, 25, 34, 43, 52, 61, and 70 ms). Furthermore, for each participant under both congruent and incongruent conditions, we separately calculated detection accuracy and fitted the data with a Boltzmann sigmoid function f(x) = 1/(1 + exp(−a × (x − x_0_))). In this function, x corresponds to the actual stimulus duration (16, 25, 34, 43, 52, 61, and 70 ms), x_0_ corresponds to the estimated stimulus duration, and a indicates the slope of the curve. We then performed paired sample t tests to compare the differences in the stimulus duration required to achieve 75% accuracy between the congruent and incongruent conditions.

#### fMRI experiment

The fMRI experiment was divided into a main experiment and a localizer experiment. For the main experiment (Figure 2a), participants were asked to focus on the odors and decide whether they smelled an odor. Each trial began with a yellow central fixation cross presented against a gray background for 1000 ms, which indicated imminent stimulus onset and signaled participants to pause their breathing and prepare to sniff. The fixation cross then changed color to blue, alerting participants to the start of a new breathing cycle. As participants inhaled, an olfactory stimulus, a visual stimulus, or both were presented for 2000 ms. There were four types of trials: unimodal olfactory (O) trials in which only the odor was presented; unimodal visual (V) trials in which only the visual image was presented at the center of the screen; bimodal congruent (OVc) trials in which olfactory and visual stimuli were presented simultaneously and carried information about the same scene, such as the smell of rose paired with a rose garden image; and bimodal incongruent (OVi) trials in which olfactory and visual stimuli were presented simultaneously but carried mismatched information, such as the smell of gasoline paired with a rose garden image. To compensate for the processing time differences between visual and olfactory stimuli, odors were delivered 700 ms before the visual stimulus. After stimulus offset, a white fixation cross appeared at the center of the screen during a variable intertrial interval randomly drawn from a pseudo exponential distribution of 8 to 12 seconds with a step size of 1 second. During the presentation of the white fixation cross, participants were asked to make a manual button press as quickly as possible to indicate whether an odor was present while they were inhaling during the presentation of the blue fixation cross. The task consisted of 5 runs, each of which included 8 trials for each of the four conditions (O, V, OVc, and OVi). Trials from the four conditions were presented in a randomized order. Before entering the MRI scanner, participants completed a practice run to ensure they were familiar with the task and experimental setup.

For the localizer experiment, participants viewed images of faces and scenes in a block design, as previously described(Epstein & Kanwisher, 1998). The functional run was composed of eighteen 12 second task blocks, including 6 blocks of rose garden scenes, 6 blocks of auto shop scenes, and 6 blocks of faces. The order of the stimulus category blocks was randomized across runs. Each block contained 15 images randomly drawn from the same category. Each image was presented for 600 ms followed by a 200 ms inter stimulus interval. Participants performed a one-back task, responding via button press whenever the same image was presented sequentially.

#### fMRI data acquisition

The MRI scanning in the current study is comprised of two data sets that were all acquired with a 3.0-T Siemens Magneton Prisma scanner, but acquired with different scan sequence. For 16 participants, the functional images were acquired using an Echo Planar Imaging (EPI) sequence (Repetition Time [TR] = 2000 ms, Echo Time [TE] = 30 ms, flip angle = 85^°^, Field of View [FOV] = 192 mm, voxel size = 2 × 2 × 2 mm) and each volume contained 32 slices. Along with functional data, a T1-weighted anatomic reference volume was acquired using a magnetization-prepared rapid gradient echo (MP - RAGE) sequence (TR = 2600 ms; TE = 3.02 ms; flip angle = 8°, voxel size = 1 × 1 × 1 mm, 176 axial slices of 1 mm thickness with no skip). For 22 participants, high-resolution functional images collected using a multislice EPI sequence (TR = 2000 ms, TE = 29 ms, flip angle = 80^°^, FOV = 192 mm, voxel size = 1.5 × 1.5 × 1.5 mm) and each volume contained 88 slices. Anatomical images acquired using a T1-weighted MP - RAGE sequence (TR = 2600 ms; TE = 3.02 ms; flip angle = 8^°^, voxel size = 1 × 1 × 1 mm, 176 axial slices of 1 mm thickness with no skip). In addition to the functional and anatomical images, a gradient echo field map images (short TE = 5.19 ms, long TE = 7.65 ms) were acquired to correct for the inhomogeneity of the magnetic field.

#### fMRI preprocessing

Standard preprocessing and statistical analyses were conducted using SPM12 (http://www.fil.ion.ucl.ac.uk/spm/). The first three volumes of each functional run were discarded to allow for saturation of the signal. After slice-timing correction, the remaining images were corrected for distortions using field map with FieldMap toolbox, realigned using the first as a reference and unwarped. Subsequently, each participant’s structural image was co-registered to the mean functional image and normalized to the Montreal Neurological Institute template brain (MNI). The transformation parameters of the structural image normalization were then applied to each EPI volume. Normalized functional data were resampled into 2-mm isotropic voxels size and were smoothed using a 6 mm Gaussian kernel for group and psychophysiological interaction analysis. And the unsmoothed EPI images in native space were used for MVPA and ROI analysis (see below).

#### Univariate analysis

At the first level, a general linear model was implemented to construct a multiple regression design matrix. Each of the four experimental conditions (O, V, OVc, OVi) were modeled as separate regressors. All events were modeled as stick functions at stimulus onset and convolved with a canonical hemodynamic response function. The six head motion parameters (3 translation and 3 rotation parameters) were included in the model as nuisance regressors. The data were high-pass filtered with a cutoff period of 128 seconds to remove low-frequency signal drifts and were corrected for serial correlations. To isolate the neural mechanisms of interest, the following contrasts were defined for each participant: (1) OVc > O and OVc > V to identify regions showing enhanced activation during congruent bimodal stimulation compared to unimodal conditions; (2) OVc > OVi to isolate regions sensitive to olfactory-visual congruency; (3) O > 0 and V > 0 to serve as inputs for conjunction analysis. The first-level contrast images were then entered into a second-level random-effects group analysis. Unless stated otherwise, regions of activation were considered significant only if they passed a cluster-level threshold of p < 0.05, family-wise error (FWE) corrected for multiple comparisons, with an underlying voxel-wise threshold of p < 0.005 uncorrected.

Subsequently, a conjunction analysis was applied to identify brain regions exhibiting olfactory-visual scene integration. The statistical criterion used to determine these regions was the intersection of four individually thresholded maps at *p* < 0.005 uncorrected. These maps included the O > 0, V > 0, OVc > V, OVc > O. The intersection of the unimodal contrasts (O > 0 and V > 0) was used to exclude regions of global deactivation and isolate regions responsive to both unimodal olfactory and visual stimuli. The remaining intersection criteria (OVc > V and OVc > O) were used to identify regions where the response to the multimodal condition exceeded the responses to either unimodal condition. To complement the statistical analysis, we also computed the percent signal changes of the resulting regions for each participant and condition. This was accomplished through a region-of-interest analysis using the MarsBaR toolbox (http://marsbar.sourceforge.net/). The regions of interest were created individually for each participant as 4-mm-radius spheres centered at the local maxima within the clusters of general linear model activations. We then extracted the mean parameter estimates from models fit to the unsmoothed data in native space.

#### Functional connectivity analysis

Given that the parahippocampal place area (PPA) and bilateral orbitofrontal cortex (OFC) were found to play critical roles in olfactory-visual scene integration, we aimed to further explore whether and how these regions interact with other brain areas involved in this process. Psychophysiological interaction (PPI) analyses were performed to identify brain regions that showed increased functional connectivity with the PPA and bilateral OFC during the processing of congruent bimodal trials relative to incongruent bimodal trials. Seed regions consisting of 4-mm-radius spheres were centered on the PPA coordinates identified in the OVc > OVi contrast, as well as on the OFC coordinates identified in the conjunction analysis. All seed regions were defined individually for each participant, and separate analyses were conducted for the PPA, left OFC, and right OFC seeds.

For each analysis, the time course of the BOLD signal was extracted from the seed region to serve as the physiological variable. The psychological variable was defined as a contrast vector representing the task effect of the congruent bimodal condition versus the incongruent bimodal condition. The PPI term was created by multiplying the deconvolved physiological variable with the psychological variable. A new first-level general linear model was then constructed that included the physiological variable, the psychological variable, the PPI term, and the six motion parameters as regressors. We specifically identified brain regions whose activation was predicted by the PPI term, which reflects a change in functional coupling with the seed region as a function of the experimental condition. The seed region’s own activity and the psychological regressor were treated as nuisance regressors. Finally, the individual contrast images for the PPI parameter estimates were entered into a second-level random-effects analysis to allow for inference at the group level.

#### Multivoxel pattern analysis

Multivoxel pattern analysis was performed using a linear support vector machine with a cost parameter of 1 as the classifier, implemented via The Decoding Toolbox (Hebart et al., 2015). This approach was used to search for brain regions whose activity could distinguish between the two categories of odors or scene images, and to determine whether classification accuracy differed across experimental conditions. For the classification samples, we extracted single-trial beta images from a first-level participant-specific general linear model that separately modeled the two odors and the two scene images. Specifically, a separate stimulus event regressor was defined for each trial in the design matrix and convolved with a canonical hemodynamic response function. Six head motion parameters were included as nuisance regressors to account for motion-related artifacts. Through this procedure, the beta parameter estimates for each trial, defined by its respective regressor within the same design matrix, corresponded to the neural response elicited by each of the 120 odor or visual trials. The resulting beta images for each trial were then entered into the classification analysis as distinct samples.

Whole-brain searchlight analyses were first conducted to identify brain regions that encoded odor or scene categories. A searchlight sphere with a 6-voxel radius was centered on each voxel to extract local activity patterns. For each participant, a whole-brain gray matter mask was employed to constrain the search space. Classification accuracy was estimated using a leave-one-run-out cross-validation procedure. In each iteration, one of the five runs served as the test set, while the remaining four runs were used as the training set. This procedure was repeated iteratively until every run had served as the test set. Binary classifications were performed separately for rose versus gasoline odors and for rose garden versus auto shop scenes. Notably, the data used for classifying rose versus gasoline were pooled from the unimodal olfactory condition, the congruent bimodal condition, and the incongruent bimodal condition. Similarly, the data for classifying rose garden versus auto shop scenes were pooled from the unimodal visual condition, the congruent bimodal condition, and the incongruent bimodal condition. Trials were collapsed across all conditions containing the relevant sensory stimuli to increase statistical power and to identify regions of interest that reliably encode task-relevant information. Consequently, a whole-brain accuracy image representing the decoding performance for the two stimulus categories was generated separately for the olfactory and visual modalities. These individual accuracy images were spatially normalized and smoothed using a 6 mm full-width at half-maximum Gaussian kernel before being entered into the group-level analysis. At the group level, a random-effects analysis was performed using voxel-wise one-sample t-tests comparing classification accuracies against the chance level of 50%. The resulting group statistical t-maps were thresholded at a voxel-level threshold of *p* < 0.005, combined with a cluster-level FEW-corrected threshold of *p* < 0.05.

The searchlight analyses served not only to replicate previous studies identifying brain regions that encoded olfactory and visual information but also to functionally define regions of interest for our main analyses. These predefined regions of interest were subsequently used in ROI-based classification analyses to track changes in neural representations across different experimental conditions. To create individual ROIs, we first manually delineated the piriform cortex on each participant’s T1-weighted anatomical image using a human brain atlas as a reference. The anatomical delineation of the orbitofrontal cortex was based on an olfactory fMRI meta-analysis(Gottfried & Zald, 2005). Scene-selective regions, including the occipital place area (OPA), parahippocampal place area (PPA), and retrosplenial cortex (RSC), were defined for each participant using data from the independent localizer runs. Following the method of Epstein and Kanwisher (1998), these regions were identified as areas responding more strongly to scenes than to faces at a threshold of p < 0.0001 uncorrected. It should be noted that because the scan sequence for the first dataset did not fully cover the anatomical location of the occipital place area, only participants from the second dataset (n = 19) were included in the occipital place area analysis.

Subsequently, voxels within the piriform cortex were functionally ranked based on their classification accuracy from the olfactory decoding map. Similarly, voxels within the PPA, OPA, and RSC were ranked according to their classification accuracy from the visual decoding map. For these four regions of interest, voxels exhibiting classification accuracy exceeding the chance level of 50% were selected to serve as independent functional ROIs for subsequent analyses. To examine changes in neural representations across different conditions, classification accuracy was estimated using a leave-one-trial-out cross-validation approach. For each of the 120 trials (comprising 60 rose and 60 gasoline trials, or 60 rose garden and 60 auto shop trials), a classifier was trained on 118 trials and tested on the remaining two trials. This procedure was repeated to cover all possible trial pairs. Finally, we calculated the mean classification accuracy separately for the unimodal olfactory condition, the congruent bimodal condition, and the incongruent bimodal condition for odors, as well as for the corresponding visual conditions.

## Acknowledgements

This work was supported by Brain Science and Brain--like Intelligence Technology – National Science and Technology Major Project 2021ZD0204200, the National Natural Science Foundation of China Grant 32430043, and the Fundamental Research Funds for the Central Universities E4EQ5001X2.

## Data, code, and materials availability

The data and code that support the findings of this study will be publicly available upon publication.

